# Identification of cytokine-predominant immunosuppressive class and prognostic risk signatures in glioma

**DOI:** 10.1101/2023.05.09.539946

**Authors:** Ziyue Tian, Zhongyi Yang, Meng Jin, Ran Ding, Yuhan Wang, Yuying Chai, Jinpu Wu, Miao Yang, Weimin Zhao

**Author notes:** Corresponding author: Weimin Zhao,.

## Abstract

**Purpose:** The advent of immune checkpoint blockade (ICB) therapies this year has changed the way glioblastoma (GBM) is treated. Meanwhile, some patients with strong PD-L1 expression remain immune checkpoint resistant. To better understand the molecular processes that influence the immune environment, there is an urgent need to characterize the immunosuppressive tumor microenvironment and identify biomarkers to predict patient survival outcomes.

**Patients and methods:** Our study analyzed RNA-sequencing data from 178 GBM samples. Their unique gene expression patterns in the tumor microenvironment were analyzed by an unsupervised clustering algorithm. Through these expression patterns, a panel of T-cell exhaustion signatures, immunosuppressive cells, and clinical features correlates with immunotherapy response. The presence or absence of immune status and prognostic signatures was then validated with the test dataset.

**Results:** 38.2% of GBM patients showed increased expression of anti-inflammatory cytokines, significant enrichment of T cell exhaustion signals, higher proportion of immunosuppressive cells (macrophages and CD4 regulatory T cells) and nine inhibitory checkpoints (CTLA4, PDCD1, LAG3, BTLA, TIGIT, HAVCR2, IDO1, SIGLEC7, and VISTA). The immunodepleted class (IDC) was used to classify these immunocompromised individuals. Despite the high density of tumor-infiltrating lymphocytes shown by IDC, such patients have a poor prognosis. Although PD-L1 was highly expressed in IDC, it suggested that there might be ICB resistance. There are many IDC predictive signatures to discover.

**Conclusion:** PD-1 is strongly expressed in a novel immunosuppressive class of GBM, but this cluster may be resistant to ICB therapy. A comprehensive description of this drug-resistant tumor microenvironment could provide new insights into drug resistance mechanisms and improved immunotherapy techniques.

## Introduction

The most frequent and severe brain tumor in adults, glioblastoma (GBM), occurs in 6 per 100,000 gliomas of the central nervous system (CNS). Additionally, there is a significant prevalence of GBM (3 cases per 100,000 people).^1^ Gliomas can be categorized into astrocytomas, ependymomas, and oligodendrogliomas based on the progenitor cell categorization. Among these, primary GBM is more common in older people and has a worse prognosis than secondary GBM. Younger people are more prone to develop secondary GBM, which results from grade II and III astrocytomas, oligodendrogliomas, or oligoastrocytomas. Unfortunately, even with vigorous therapy, the median survival in the GBM class is just 13.5 months despite advances in the understanding of the genesis and prognosis.^2–3^

Patients with melanoma and other tumors have responded favorably to ICB treatment.^4–7^ The US Food and Drug Administration(FDA) has so far authorized 3 therapeutic vaccinations.^8^ While some patients with solid tumors have demonstrated a survival advantage with immunotherapy, the majority continue to be resistant. Less than 15% of cancer patients react to ICB at the moment.^9^ The use of immunomodulation for the treatment of GBM or patient survival has not yet significantly benefited clinical studies using ICB and vaccination therapy.^10–11^

An allogeneic IDC/autologous GBM vaccine in combination with granulocyte-macrophage colony-stimulating factor (GM-CSF), cyclophosphamide,^12^ and bevacizumab was shown to significantly improve survival in phase II clinical study, according to interim results from that study.^13^ However, administration of anti-programmed cell death protein 1 (PD-1) mAb before tumor resection increased both local and systemic antitumor immune responses The preclinical outcomes of immunomodulation in conjunction with other therapies have been favorable.

Although the very complex and varied tumor microenvironment of GBM is present, it is yet unknown how this environment may affect the effectiveness of GBM immunotherapy. In this study, we followed the methods of Minglei Yang et al. ^14^ and aimed to analyze RNAseq expression data from 278 human GBM samples and to isolate transcriptomic signals released by immunosuppressive TMEs via UCC.^15^ This was accomplished by deconvoluting the gene expression signals from tumor cells, inflammatory cells, stromal cells, and cytokines from a large number of tumor samples. This led to the identification and validation of a GBM immune type with immunosuppressive molecular characteristics and potential ICB resistance. To examine the underlying causes and survival prognosis of ICB, the depleted immune panel may also be connected with multi-omics data.

## Material and methods

### Data download and processing

Gene expression profiling data were obtained from The Cancer Genome Atlas (TCGA) (151 RNA-seq datasets) and the Chinese Glioma Genome Atlas (CGGA) (127 microarray datasets).^16–17^ Since different tumor stages lead to different clinical features and treatment options, we selected TCGA GBM patients for training and CGGA patients for validation. UCC analysis was performed on data from protein-coding genes.

### Identifcation of depletion immune class by unsupervised Clustering

Consensus clustering is a technique for unsupervised clustering that uses gene expression data to identify unidentified categories of illnesses or tissues.^18^ By choosing the R language’s ConsensusClusterPlus package, consensus clustering analysis is carried out. The algorithm’s parameters, including maxK, clusterAlg, and distance, are set to 6, “hc”, and “Pearson”, respectively. By monitoring the cell, the cumulative density function, and the relative change in the area under the cumulative density function, the ideal number of immunological subtypes (K) was determined. To pinpoint important subgroups connected to GBM prognosis, we used gene expression patterns from the TCGA database linked with TCGA-GBM.

Using single-sample gene set enrichment analysis (ssGSEA), immunological and stromal enrichment scores were generated by other studies^19–21^ to identify expression patterns relevant to the immune and stromal systems.^22^ We saw that cluster 1 had a higher enrichment score compared to the other clusters when the enrichment scores from the immunological and matrix enrichment analyses were merged with the 2 UCC discovered groupings. As a result, Cluster 1 is referred to as the “Immune stromal Clusterℍ throughout this text.

Next, a study examined the prevalence of certain immune cells in cancers associated with the immune stromal cluster. The expression profiles of ssGSEA were used to determine enrichment scores after collecting signatures for various immune cells. To ascertain the prevalence of certain immune cells in the immune stromal clusters, the enrichment fractions of 28 immune cells were merged with the clusters^23^. Additionally, the CIBERSORT algorithm’s inferences of the absolute fractions of 22 invading immune cells based on gene expression patterns were obtained from the TIMER database.^24^ Leukocyte fraction data from the Torsson trial, determined by DNA methylation, from https://gdc.cancer.gov/about-data/publications/panimmune.25 The Saltz study’s Supplementary Table lists the leukocyte fraction determined by pathological imaging of TCGA tumors, including GBM.^26^ The immune stromal class was compared to the remaining clusters to verify the enrichment of lymphocytes in the Immune stromal clusters.

Finally, the differential expression of nine immunosuppressive checkpoints (CTLA4, PDCD1, LAG3, BTLA, TIGIT, HAVCR2, IDO1, SIGLEC7, and VISTA) was analyzed.

Differential genes with two expression patterns were identified by differential analysis.

Enrichment analysis of differential genes to study biological processes with significant differences in two expression patterns.

### Validation of IDC in CGGA to confrm the presence of depletion immune

Using the same technique as above, we ran UCC and ssGSEA analysis on the RNAseq-based expression profiles of 127 CGGA samples to validate the occurrence of an immune depletion state in GBM. Similarly, we observe 2 GBM clusters. We noticed that Cluster 2 had higher enrichment scores for immune cell, stromal, and T cell exhaustion (tex) associated signatures. As a result, Cluster 2 was determined to be the GBM IDC. Between the IDC group and other groups, the ratios of immune cells, white blood cells, TIL, and the expression of numerous inhibitory receptors were compared. The analyses by GSEA were conducted to evaluate the enrichment of molecular pathways and gene expression signatures in the IDC. The hallmark gene sets were collated from MSigDB (https://www.gsea-msigdb.Org/gsea/msigdb).

#### Drug sensitivity analysis

Cancer Drug Sensitivity Genomics was used to gather data on drug responsiveness and drug targeting pathways (GDSC). The pRRophetic package in the R language was then used to predict the drug sensitivity of various phenotypes from the gene expression data and obtain the drugs associated with the various classes after counting the sensitivity data of the two GBM classes to various drugs. Establish a foundation for choosing clinical drugs.

### Establishment of clinically applicable prognostic models

Significant genes associated with survival were then found using univariate COX regression analysis and least absolute shrinkage and selection operator (LASSO) regression, and a risk prediction model was created. The formula for calculating the risk score is as follows:

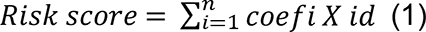

where X is the normalized count for each gene and Coefi is the coefficient. We classified patients into high-risk and low-risk groups based on the median risk score. Both the training set and the test set were made up of follow-up data from GBM data in the TCGA and CGGA, respectively. By using survival curves from the training and test sets, within-group validation was used to evaluate the reliability of the risk scoring model. Based on the training and testing sets, the survival rates of patients at 1, 3, and 5 years were projected. A forest plot of clinical predictive indicators was created by combining risk scores and clinical traits to acquire clinical data regarding GBM survival and prognosis.

### Prognostic Nomogram Construction

For further exploring the prognostic role of clusters, univariate and multivariate regression analysis was also performed to filter the significant prognostic factors. To determine which clusters were significantly associated with OS, the univariate regression models were used and p-values were calculated. The gene classifier signatures with a p-value <0.05 are significantly related to clinicopathologic features. Subsequently, a prognostic nomogram was constructed to improve the prediction ability by integrating important features derived from multiple regression analysis with the survival and RMS packages. The calibration plot, a common chart used to evaluate the consistency of the nomogram, was gen-erated by using the rms package of R software.

### Gene set variation analysis

GSVA (gene set variation analysis) is a non-parametric, unsupervised algorithm that can calculate the enrichment score of a specific gene set in each sample. Immune depletion scores were calculated for the two subtypes by using the risk model as the immune depletion label, and the immune depletion score was calculated by Wilcox test with the expression of nine immunosuppressive checkpoints (CTLA4, PDCD1, LAG3, BTLA, TIGIT, HAVCR2, IDO1, SIGLEC7, and VISTA)

### Correlations between IDC and rest class PD-1 and TGF-β expression and the ICB response

Patient ICB treatment responses were predicted using the Tumour Immune Dysfunction and Exclusion (TIDE) algorithm. To investigate IDC patient responses to ICB treatment, programmed death-ligand 1 (PD-L1) expression was compared between IDC and resting classes. Higher TIDE prediction scores were generally associated with worse ICB responses.

According to Mariathasan et al., the cytokine transforming growth factor-β (TGF-β encoded by TGFB1) inhibits antitumour immunotherapies.^27^ By contrast, therapeutic co-administration of anti-TGF-β blockade-inducing antibody and anti-PD-L1 antibody reduced TGF-β signaling in stromal cells, promoted T cell infiltration into the tumour centre and stimulated strong antitumour immunity and tumour regression. Taken together these observations suggest that TGF-β acts by limiting T cell infiltration of the TME to thereby suppress antitumour immunity. Therefore, detection of TGF-β in TNBC patient tumours correlates with resistance to antitumour immunotherapies.

### Reverse transcriptase-PCR Analysis

Human astrocyte cell line (HA1800) and human GBM cell line (U118) were purchased from Shangcheng North Na Chuanglian Biotechnology Co., LTD. The expressions of PDL1 and TGFB1 in GBM cell lines and human astrocyte cell lines were analyzed in the control group of HA1800 and U118. Total RNA was extracted using the Redzol kit from Beijing SBS Gene Technology Co., Ltd., and was extracted according to the instructions.

The forward primer sequence is TGFB1: F-5′-CCTTGCTGGACCGCAACAAC-3′, the reverse primer sequence is TGFB1: R-5′-CAGCAGCCGGTTACCGAC-3′, and the product length is 153bp; PDL1: F-5′-TTTGCTGAACGCCCCATACA-3′, the reverse primer sequence is PDL1: R-5′-TGTCCCGTTCCAACACTGAG-3′ and the product length is 264bp. qRT-PCR was performed using the SYBR® Premix Ex Taq™ II (Takara, Shiga, Japan) Kit and a StepOnePlus Real-Time PCR instrument. Briefly, the mixture contained SureScript RTase Mix (20×) 1 μL, SureScript RT Reaction Buffer (5×) 4.0 μL, Total RNA 1 μg and dd HO (RNase/DNase free) supplemented to 20 μL, then sealed the cDNA with a transparent sealing film Array, mix well, spin off for 5s, 5 min at 25°C, 15 min at 42°C, 5 min at 85°C, keep at 4°C, and store the product at -20°C. qRT-PCR reactions were performed on a LightCycler 96 Real-Time PCR System (Roche Diagnostics, Indianapolis, IN). The reaction mixture was activated at 50°C for 2 min, pre-denatured at 95°C for 10 min, and then subjected to 40 cycles of amplification reactions at 95°C for 15 s and 60°C for 30 s. Finally, LightCycler 96 software (version 1.1.0.1320, Roche) was used for the collection and analysis of qRT-PCR data. With β-actin as an internal reference gene, the relative expression of mRNA was calculated by 2^-ΔΔCt^ method.

### Statistical analysis

R (version 4.2.1, http://www.rproject.org) was used for statistical discrete analysis. The Wilcoxon rank sum test for continuous data was used to correlate IDC and the remaining categories with tumour-infiltrating lymphocyte (TIL) percentage and neoantigen number. Overall survival (OS) data were analysed using Kaplan-Meier estimates and log-rank testing. To discover variable combinations, we included all clinicopathological factors in the Cox model. P-values of ≤0.05 were considered statistically significant. Pearson’s correlation was used to assess the strengths of two-variable linear relationships. Genomic mutation data of all CGGA tumour samples were obtained from the TCGA database.

## Results

### Identifcation and characterization of a novel depletion immune class in GBM

The training cohort’s 151 GBM samples’ RNAseq-based gene expression profiles underwent extensive UCC analysis, and TEX-related transcriptomic signals were extracted in the TME. Two expression clusters were robustly created from the training cohort’s dataset (Figure 1A). A considerable enrichment of immune cell and stromal component gene expression profiles was shown by the high immune and stromal enrichment fractions computed by ssGSEA and batch RNAseq-based gene expression profiling in GBM patients in cluster 1. This group was subsequently given the moniker Immune Stromal Cluster (Figures 1B-C). Immune cell subsets, T cells, neutrophils, monocytes, and other immune cell signatures were also evident in the immune stromal clusters (Figures 1D). The Immune Stromal Cluster’s enrichment of immune cells was further confirmed, and CIBERSORT’s estimation of the absolute proportion of immune cells in the Immune Stromal Cluster compared to the other clusters using batch RNAseq data. The proportions of CD4 T cells, monocytes, and neutrophils in the immune stromal clusters were greater than those in other clusters, supporting the enrichment analysis carried out using ssGSEA (Figure 1D). Additionally, compared to the other clusters, the Immune stromal cluster had a considerably larger leukocyte proportion calculated by DNA methylation (Figure 1E). Similarly, the Immune stromal cluster has much more TILs than the other clusters as determined by pathological imaging. Several inhibitory receptors, including CTLA4, PDCD1 (also referred to as PD-1), LAG3, BTLA, TIGIT, HAVCR2 (also referred to as TIM-3), IDO1, SIGLEC7, and VISTA, were profiled for their expression levels in order to investigate TEX in GBM (Figure 1F). A novel subclass inside the Immune stromal cluster, which comprises 38.2% of the training cohort and is henceforth referred to as the immune depleted class (IDC), was discovered based on the above-described expression analysis of inhibitory receptors and enrichment scores for the TEX signaling gene collection. Through differential analysis, 311 genes were found to be differentially expressed between the two classes, among which 132 genes were down-regulated and 179 genes were up-regulated (Figure 1G, Supplementary file 1). In the GO enrichment results of these differential genes, they are mainly enriched in humoral immune response, adaptive immune response based on somatic recombination of immune receptors built from immunoglobulin superfamily domains and leukocyte migration(Figure 1H).

**Figure 1.**
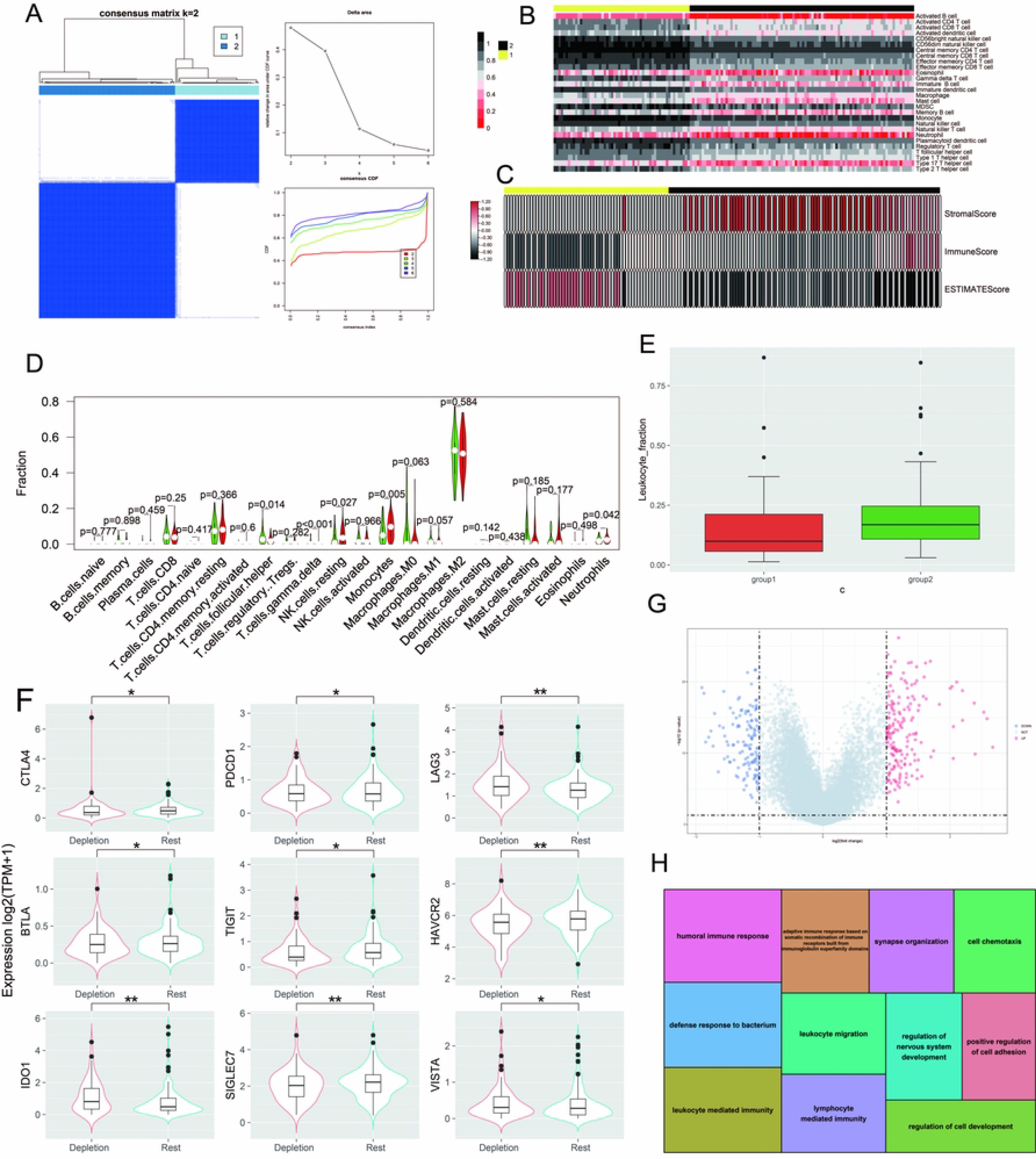
The identifcation and molecular characterization of IDC. (A) Consensus clustering for the TCGA-GBM. (B) Stromal and immune enrichment analysis defined the cluster 2 of two expression patterns as an immune-stromal cluster. High and low gene enrichment scores are delineated in black and red, respectively. (C) The enrichment scores of gene signatures identifed the immune cells for the immune-stromal and other clusters. (D) The comparison of the absolute fractions of TME cells inferred by CIBERSORT. (E) Box plots show the diferences of leukocyte fraction and percentage between two classes. (F) Box plots show diferent expression levels of multiple inhibitory receptors in the immune-stromal cluster compared to the other clusters. (G) Volcano plot of difference analysis. (H) Enrichment analysis plot. *: P < 0.05; **: P < 0.01.

### Validation of IDC in TCGA GBM

We use UCC analysis on bulk RNAseq-based gene expression profiles of CGGA samples to confirm the presence of reduced immune classes in GBM and subsequently got 2 clusters (Figure 2A-B). We noticed that cluster 2 had greater stromal and immune enrichment scores than the other clusters when the feature enrichment scores generated by ssGSEA with bulk RNAseq-based gene expression profiles were combined with the clusters (Figure 2D). Features and other cytokine indicators linked to TEX were considerably enhanced (Figure 2C). As a result, the depleted Immune class in the CGGA sample is classified as Cluster 2. In the IDC of CGGA, numerous inhibitory receptors were co-upregulated (Figure 2E). GSEA enrichment results are mainly enriched in HALLMARK_INFLAMMATORY_RESPONSE, HALLMARK_IL6_JAK_STAT3_SIGNALING and HALLMARK_INFLAMMATORY_RESPONSE (Figure 2F).

**Figure 2.**
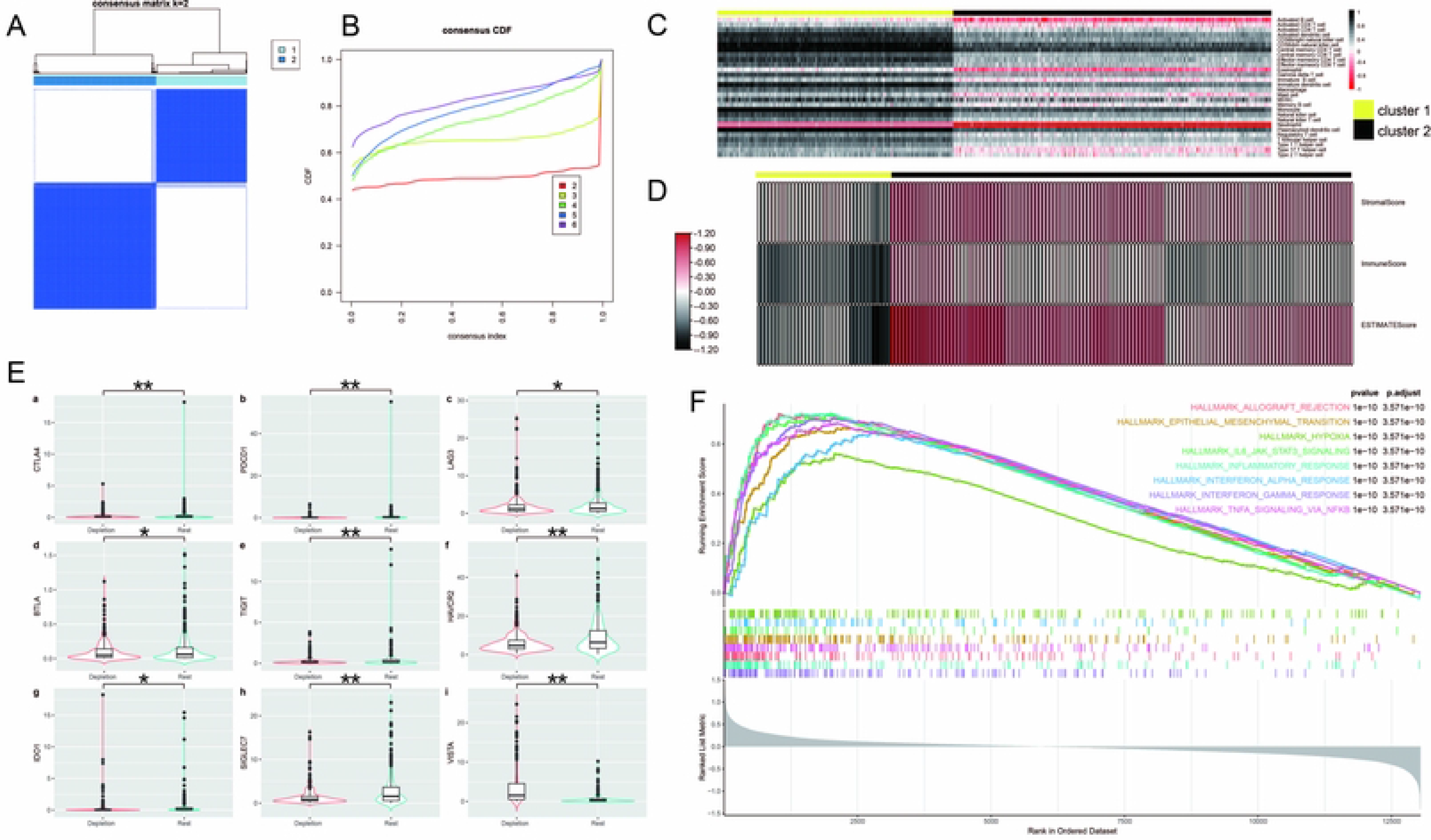
Internal validation of IDC in CGGA samples. (A-B) Consensus clustering for the CGGA. (C) Heatmap shows the cluster calculation of the enrichment of immune cells by ssgsea. (D) The enrichment scores of gene signatures identifed the immune cells for the IDC and rest clusters. (E) Boxplots shows the diferent expression levels of multiple inhibitory receptors between two classes. (F) GSEA analysis plot. *: P < 0.05; **: P < 0.01.

### IDC had poorer prognosis in GBM

According to Kaplan-Meier assessment, the OS of IDC was significantly lower than that of the rest class overall (P < 0.001; Figures 3A), and the OS of IDC patients with early GBM was significantly lower than that of the rest class (P < 0.001; Figures 3B). OS in advanced stage was also lower in IDC patients than in the rest class (P < 0.001; Figures 3C-D). These results suggest that although T cells are widely distributed in IDCs, most of them are immunosuppressed, losing the ability to stop tumor growth and leading to a poor prognosis.

**Figure 3.**
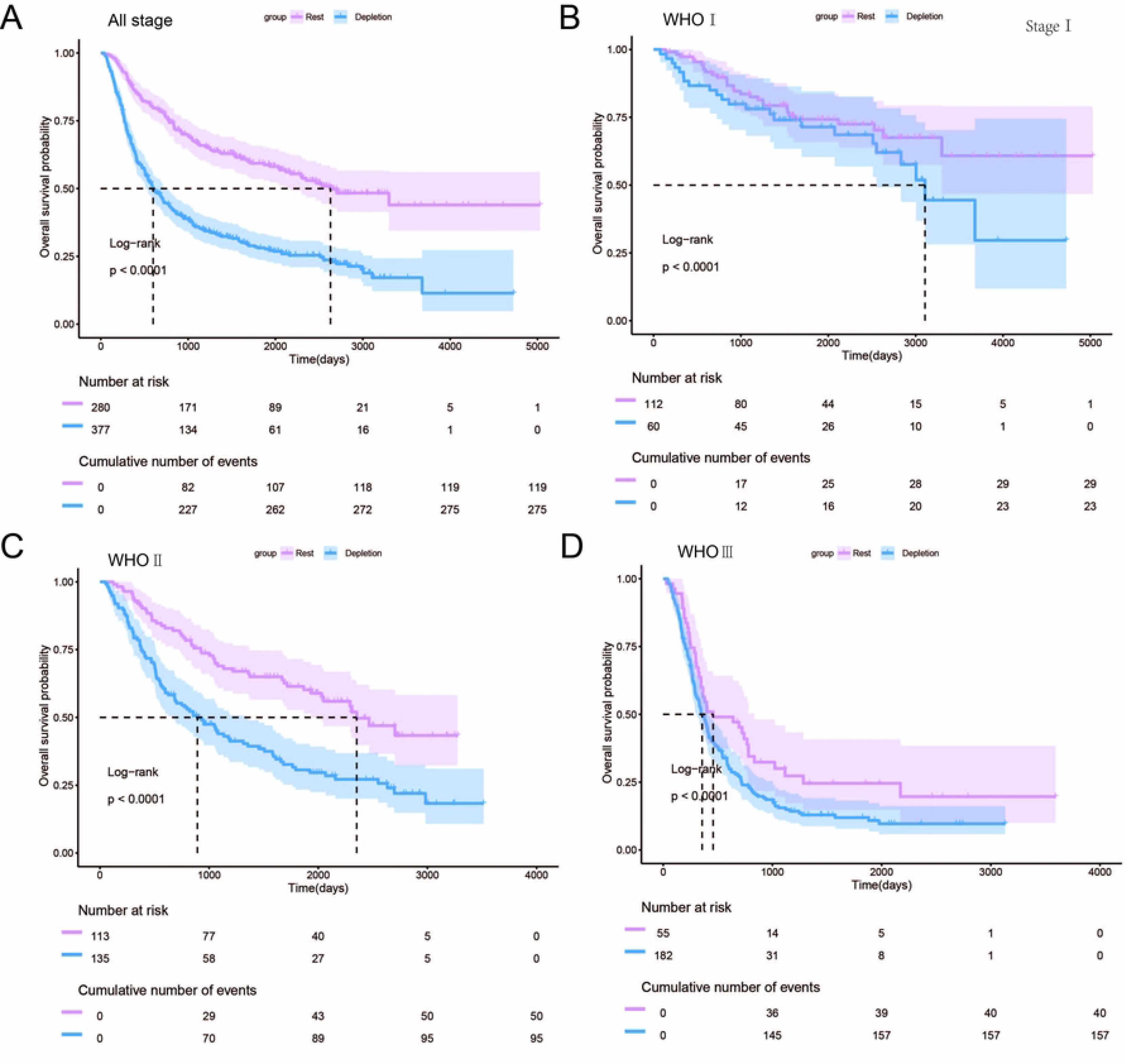
Prognosis analysis for the IDC and the rest class across diferent stages of GBM. (A-D) Kaplan Meier estimates of overall survival for the IDC and the rest class across all-stage GBM and WHOI, II, III.

### Immune depletion traits can predict how chemotherapy work

Using the “pRRophetic” package of R, the IC50 differences of eight targeted and chemotherapeutic drugs in two class of clustering were studied to predict their sensitivities to drug therapy. Figure 10a–h, respectively, shows the drug sensitivity results of Sorafenib, Gefitinib, Bleomycin, Bosutinib, Etoposide, Lenalidomide, Camptothecin, and Methotrexate in three subtypes of GBM class. The statistical results demonstrated that the Rest class are generally more sensitive to tumor treatment drugs than the IDC (Figures 4A-H).

**Figure 4.**
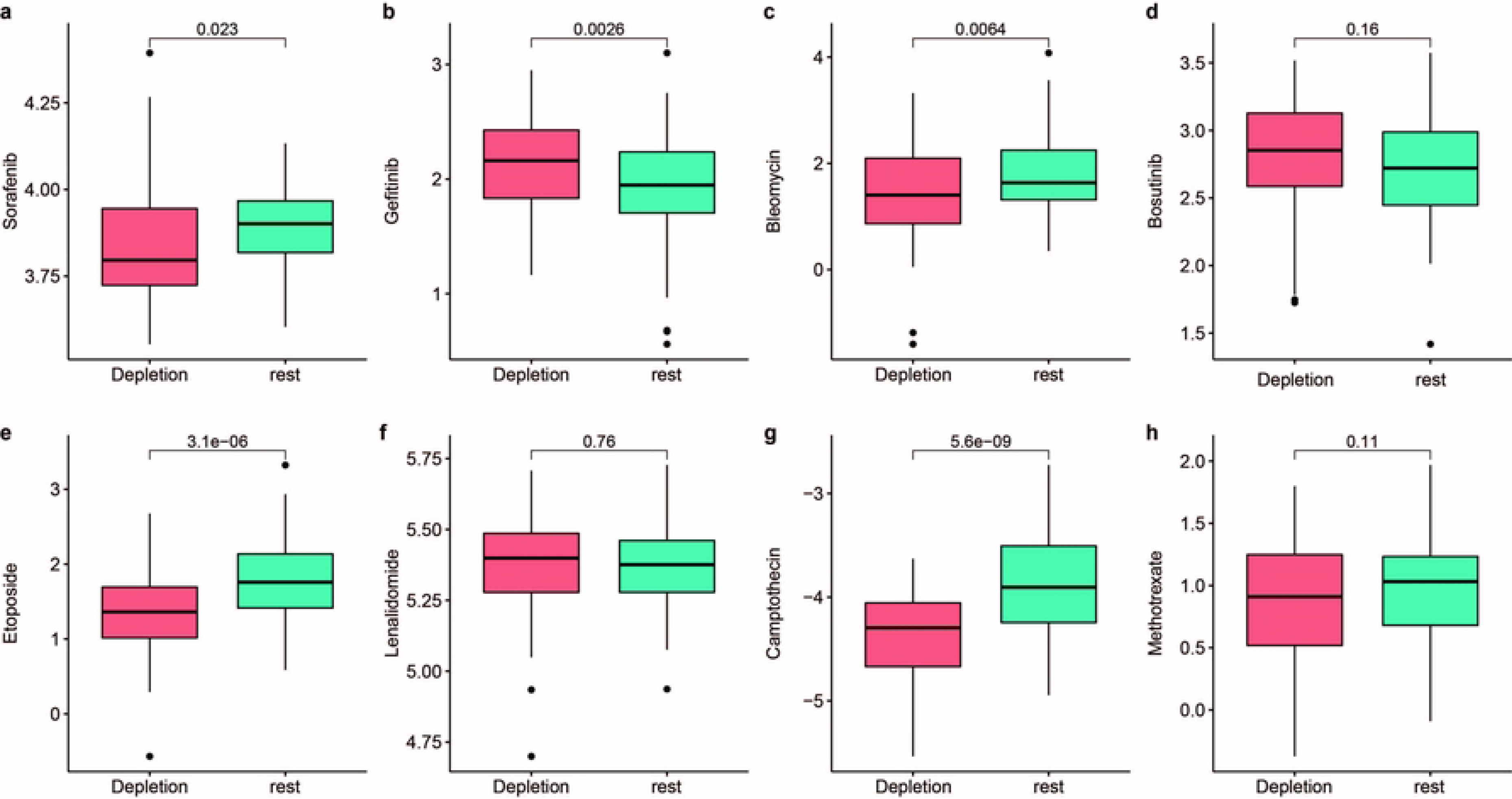
Drug sensitivity analysis. (A-H) Gene expression profile characteristics of triple-negative breast cancer patients, drug sensitivity test (Sorafenib, Gefitinib, Bleomycin, Bosutinib, Etoposide, Lenalidomide, Camptothecin, Methotrexate), analysis of drug IC50 in patients with different immune depletion conditions. A-H Drug sensitivity IC50 of patients under different immune depletion conditions. I-P Drug sensitivity IC50 of patients with high and low risk scores.

### Construction and validation of the prognostic prediction model

The univariate Cox regression algorithm was used to initially obtain 311 genes related to GBM patient prognosis then hazard ratios (HRs) and P values of these genes were calculated. Next, we constructed risk models using the LASSO algorithm that ultimately led to the identification of 26 prognostically relevant genes (Figures 5A-C). These genes were then used to construct risk score-based models based on training (n = 127) and test (n = 151) datasets obtained from TCGA-GBM and CGGA GBM patients, respectively. Results of survival analysis revealed that higher risk scores in training and test sets corresponded to poorer survival (P < 0.0001) (Figures 5 D, E, G, H). Time-dependent ROC curves were generated and used to assess the sensitivity of the prognostic model. Results obtained for the areas under ROC curves (AUCs) revealed 3-, 5- and 10-year AUCs for the training set of 0.741, 0.82 and 0.867, respectively (Figure 5F), and corresponding AUCs for the test set of 0.679, 0.688 and 0.689, respectively (Figure 5I).

**Figure 5.**
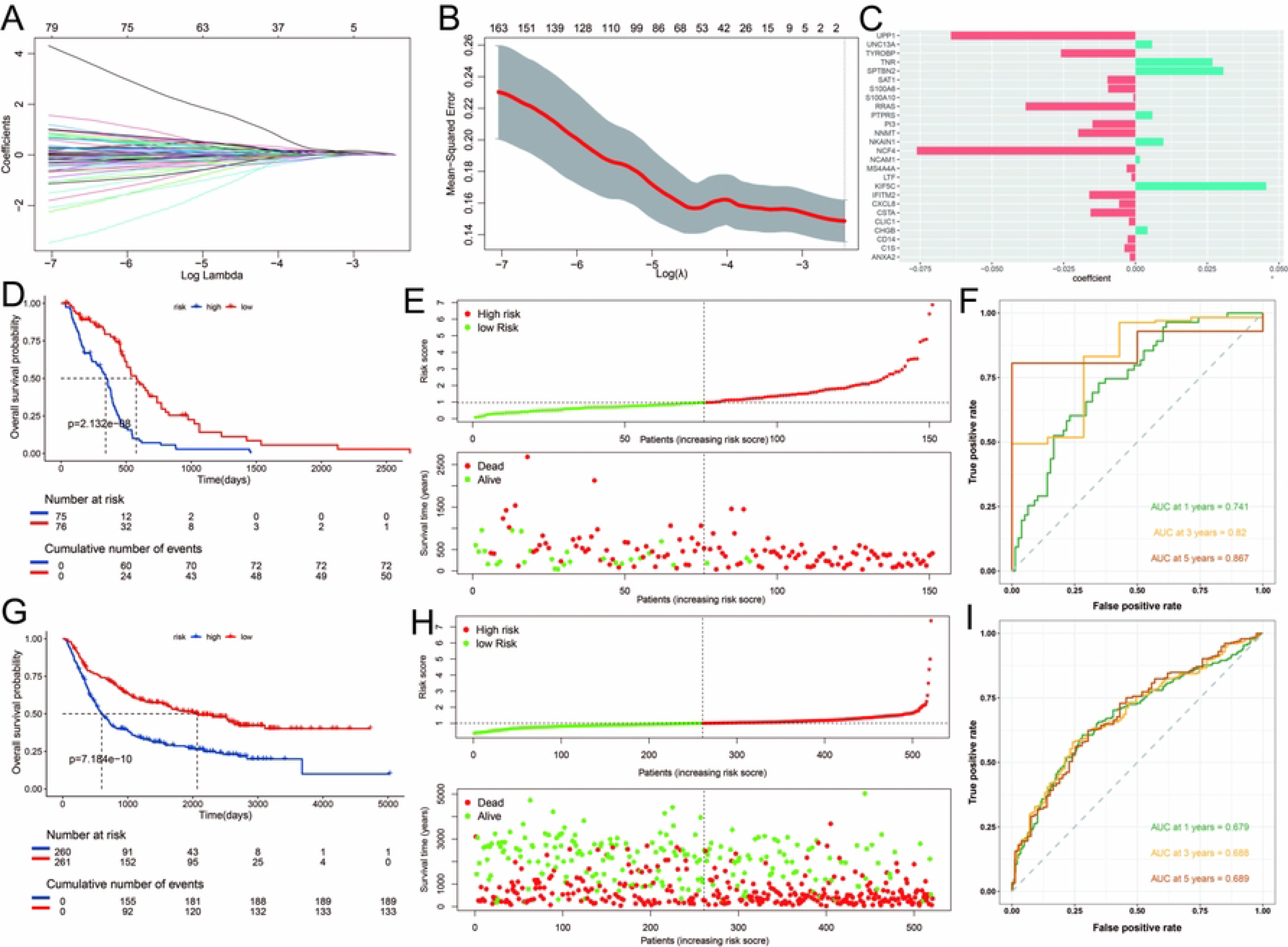
Construction of the risk model. (A) Partial likelihood deviance was revealed by the LASSO regression model in the 10-fold cross-validation. The vertical dotted lines were drawn at the optimal values by using the minimum and 1-SE criteria. (B) LASSO coefficient profiles of 22 selected genes in the 10-fold cross-validation. The vertical dotted lines were drawn at the optimal values by using the minimum criteria and 1-SE criteria. (C) The forest plot of the associations between the infiltrate levels of 26 prognostic molecules and OS in the training cohort. The HR, 95% CI and P-value were determined by univariate Cox regression analysis. (D, G) Kaplan-Meier curves for the test and training sets, respectively (P<0.001, log-rank test, survival rate comparison). (E,H) Patients were divided into high-risk and low-risk subgroups based on median level of CRRS in TCGA and CGGA; survival status of patients in two subgroups. (F, I) Time-dependent receiver operating characteristic (ROC) of training and test sets. I Proportion of IDC and rest class in high and low risk patients.

### Independent Prognostic Ability of Groups

For evaluating the independent prognostic ability of the obtained gene classifier signatures, we also investigated the correlation between the gene classifier signatures and clinicopathological features of TCGA–GBM through univariate regression analysis and multivariate Cox regression analysis. The clinically common factors like Grade, Age, IDH_mutation_status, MGMTp_methylation_status and Group were registered as covariates for analysis. As a result, Grade, IDH_mutation_status and Group are independent factors that can be used to predict the prognosis of GBM patients (Figure 6A,B). In addition, the clinicopathological information and independent prognostic factors are combined and then a nomogram involving three items in GBM was constructed, which is used as a clinically relevant quantitative method for predicting the mortality of GBM patients. Obviously, we found that by adding points for each prognostic parameter, each patient will be assigned a total score value and higher total scores corresponded to poorer patient outcomes (Figure 6C). Time-dependent C-index curves of different variables based on TCGA cohorts show the optimum performance of nomogram compared with other single factors (Figure 6D). Moreover, calibration plots for the TCGA GBM reveal a similar performance of the nomogram to the developed prognostic model (Figure 6E). The correlation between clinical factors and different classes is calculated by chi-square test, and the distribution of significantly related clinical factors in each class is displayed by histogram, there are more high-grade stages distributed in IDC, older patients and more wildtype IDH mutation (Figure 6F).

**Figure 6.**
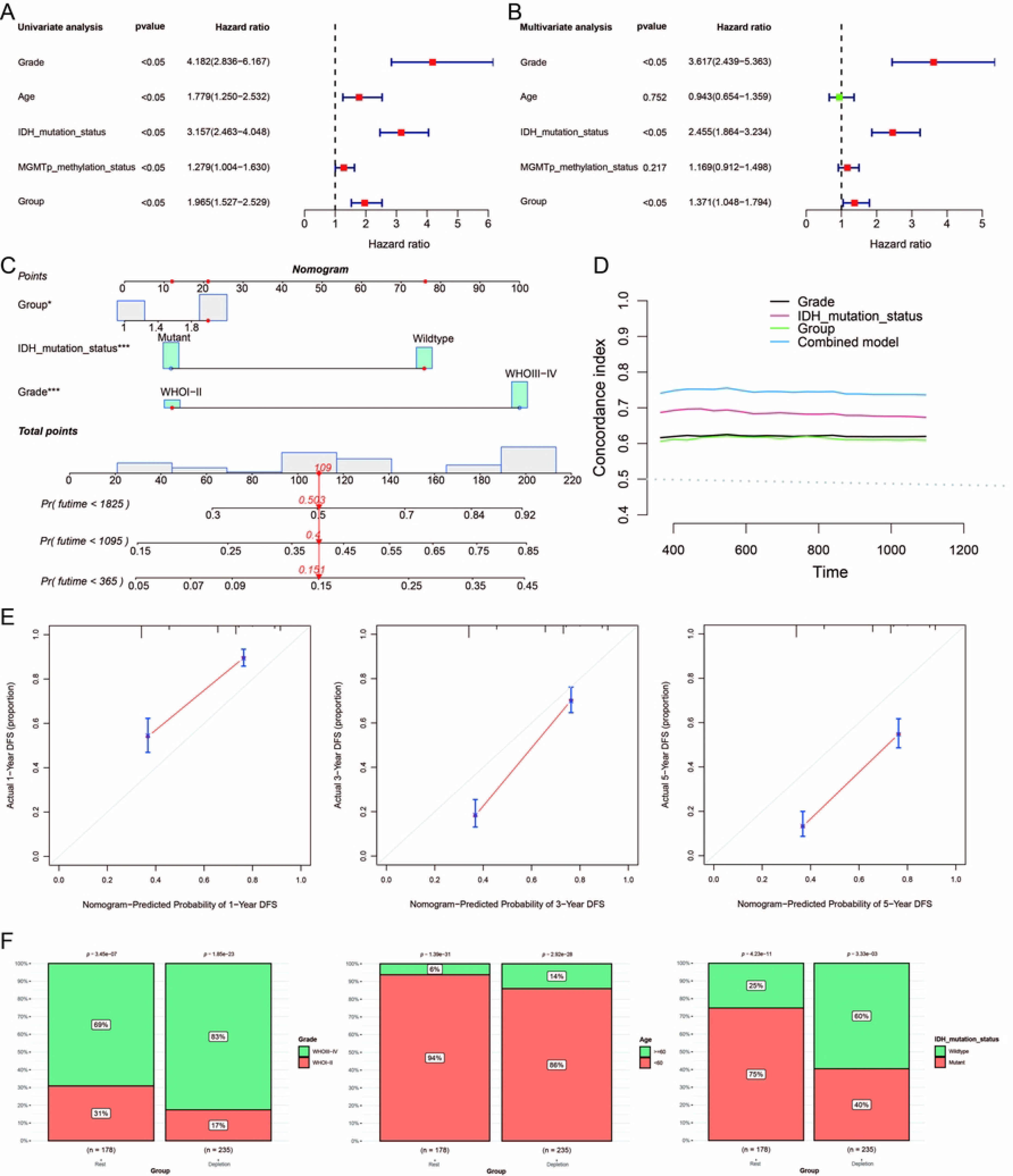
Correlation between the groups and clinicopathological features of TCGA–GBM samples. (A, B) Univariate and multivariate analysis including groups and clinical factors. (C) Nomogram for comprehensively predicting the 1-, 3-, and 5-year OS of BC patients in the TCGA database. (D) Time-dependant c-index plot for the nomogram and other clinical factors. (E) Calibration plots for estimating the 1-, 3-, and 5-year OS for TCGA–GBM samples.

### IDC scoring and immunosuppressive checkpoints analysis

By using the risk model as a gene set to perform GSVA analysis on TCGA-GBM data, the IDC score of each TCGA sample was obtained, and the IDC score of IDC and rest class were found to be significantly different through the Wilcox test, suggesting that the model is effective for clinical GBM tumors Patient has evaluative significance (Figure 7A). Next, the correlation between the expression of nine immunosuppressive checkpoints (CTLA4, PDCD1, LAG3, BTLA, TIGIT, HAVCR2, IDO1, SIGLEC7, and VISTA) and the IDC score is displayed by a scatter plot. The correlation analysis shows that the IDC score and the immunosuppressive checkpoints are positive Correlation, suggesting that the stronger the expression of immunosuppressive checkpoints in IDC, the higher the level of IDC (Figure 7B).

**Figure 7.**
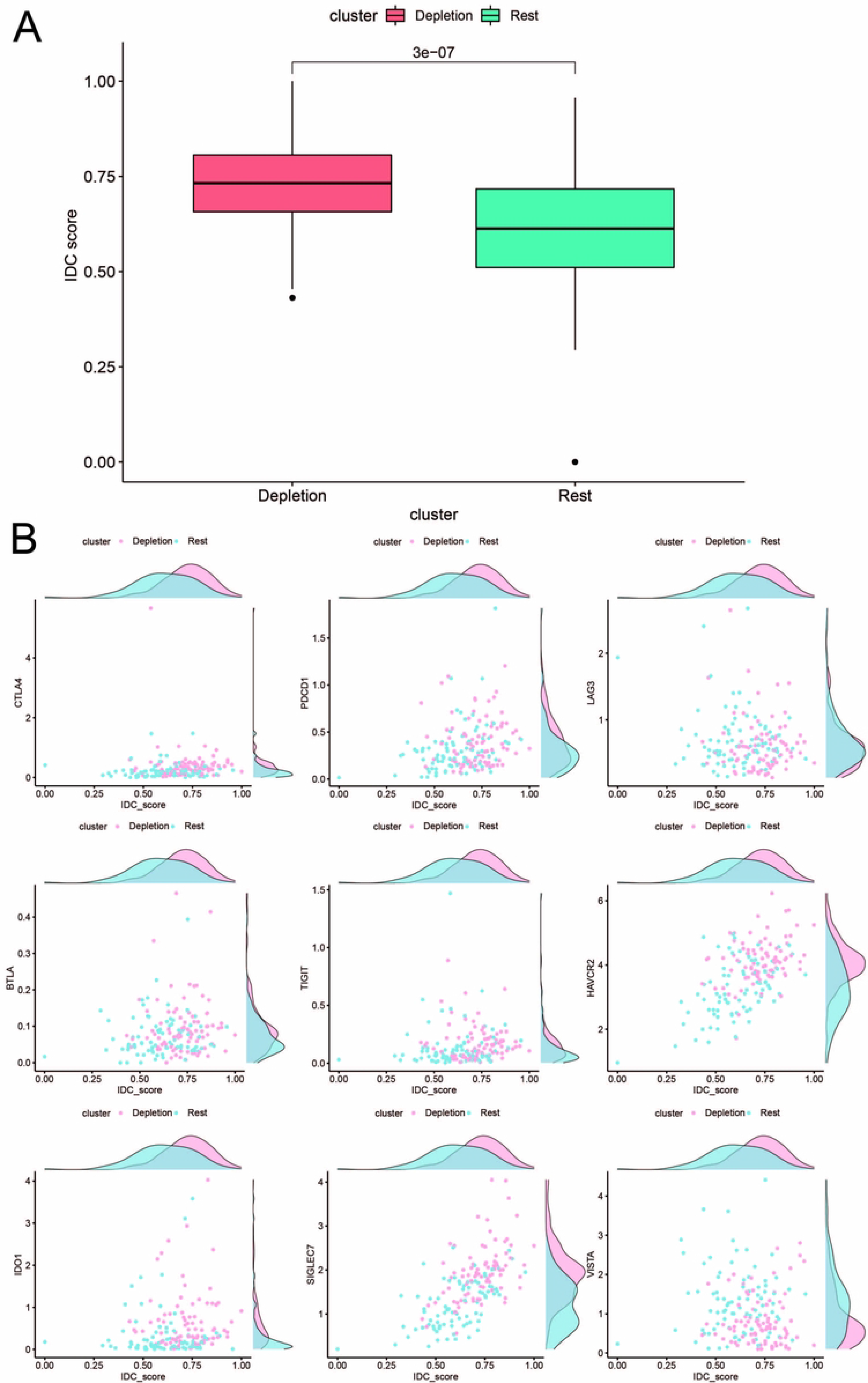
(A)The expression level of IDC score between IDC and rest class based on TCGA. (B) Correlation between IDC score and the expression of nine immunosuppressive checkpoints (CTLA4, PDCD1, LAG3, BTLA, TIGIT, HAVCR2, IDO1, SIGLEC7, and VISTA) from TCGA sample.

### The IDC immunosuppressive signature correlates with immunotherapy resistance

Treatment of cancer patients with therapeutic antibodies targeting the PD-L1 pathway can elicit long-lasting, robust responses. However, efficacies of such treatments are frequently reduced due to the emergence of drug resistance. To study the response of the IDC group to ICB treatment, we compared PD-L1 expression levels between IDC and rest patient groups and found that PD-L1 expression levels were higher in the IDC group than in the rest group (Figures 8A, B). Next, use of the Tumour Immune Dysfunction and Rejection (TIDE) algorithm to predict ICB treatment responses revealed that both early and advanced GBM IDCs had higher TIDE prediction scores than did the other categories (Figures 8C, D), such that a higher TIDE prediction score indicated a worse ICB response. Importantly, TGFβ1 was expressed at a lower level in rest classes than in IDCs (Figures 8E, F), which is consistent with results obtained by Mariathasan et al. that indicated the cytokine TGF-β (encoded by TGFβ1) suppresses effects of antitumour immunotherapy.^47^ Taken together, these results suggest that the IDC immunosuppressive signature has a ICB therapy resistance.

**Figure 8.**
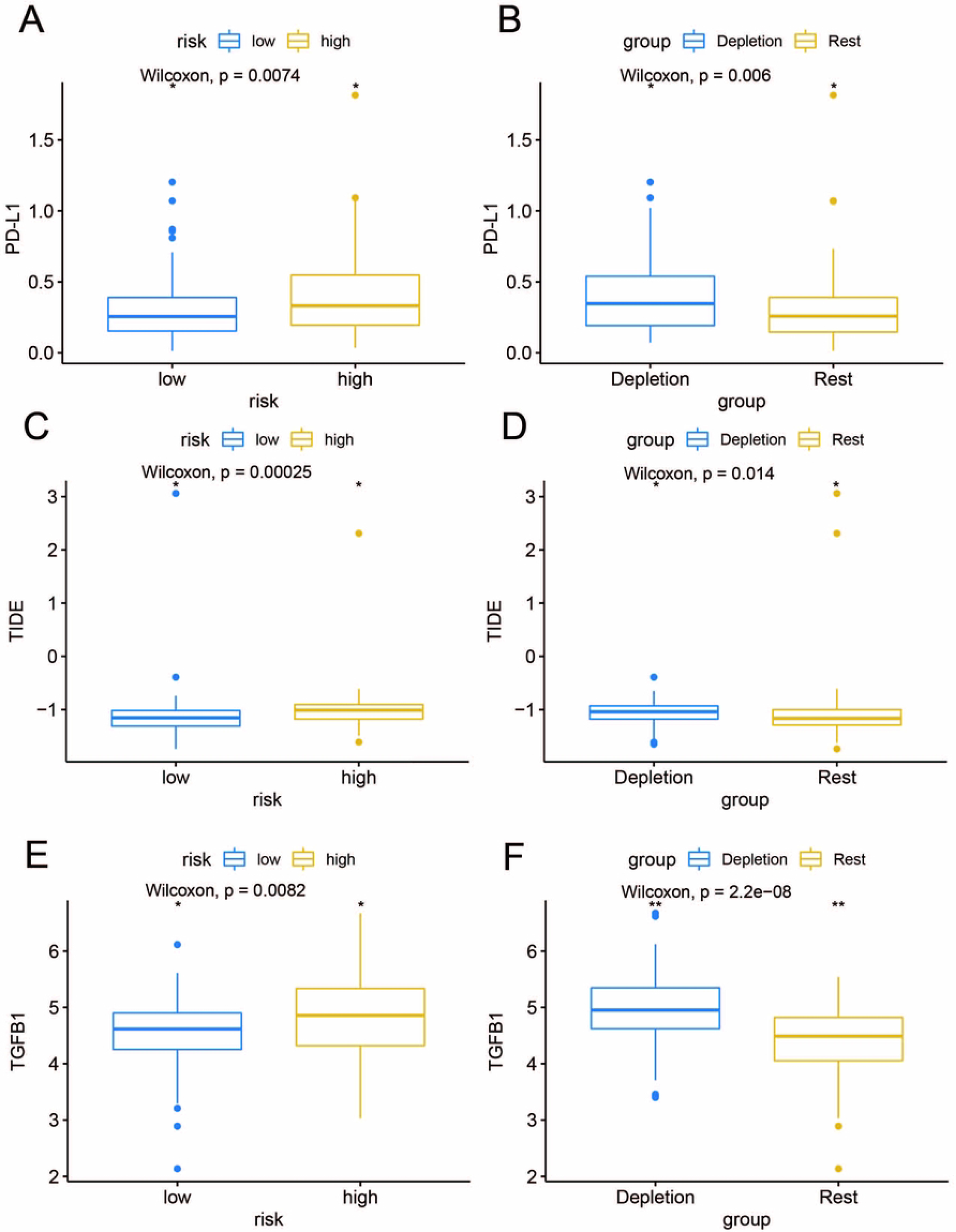
Prediction of resistance to ICB therapy. (A-B) Different expressions of PD-L1 expression levels in different classes. (C–D) Different expression of TIDE predictive score for different classes. (E-F) Different expression of TGFβ1 in different classes. *: P < 0.05; **: P < 0.01.

### GBM gene expression level verification via quantitative reverse transcription PCR (qRT-PCR)

Expression levels of selected genes were determined using qRT-PCR to confirm their reliability. As shown in Figure 9, the results for all mRNA transcripts showed that they were expressed at significantly higher levels in the GBM tumor cell lines (U118) than in the adjacent normal cell lines(HA1800).

**Figure 9.**
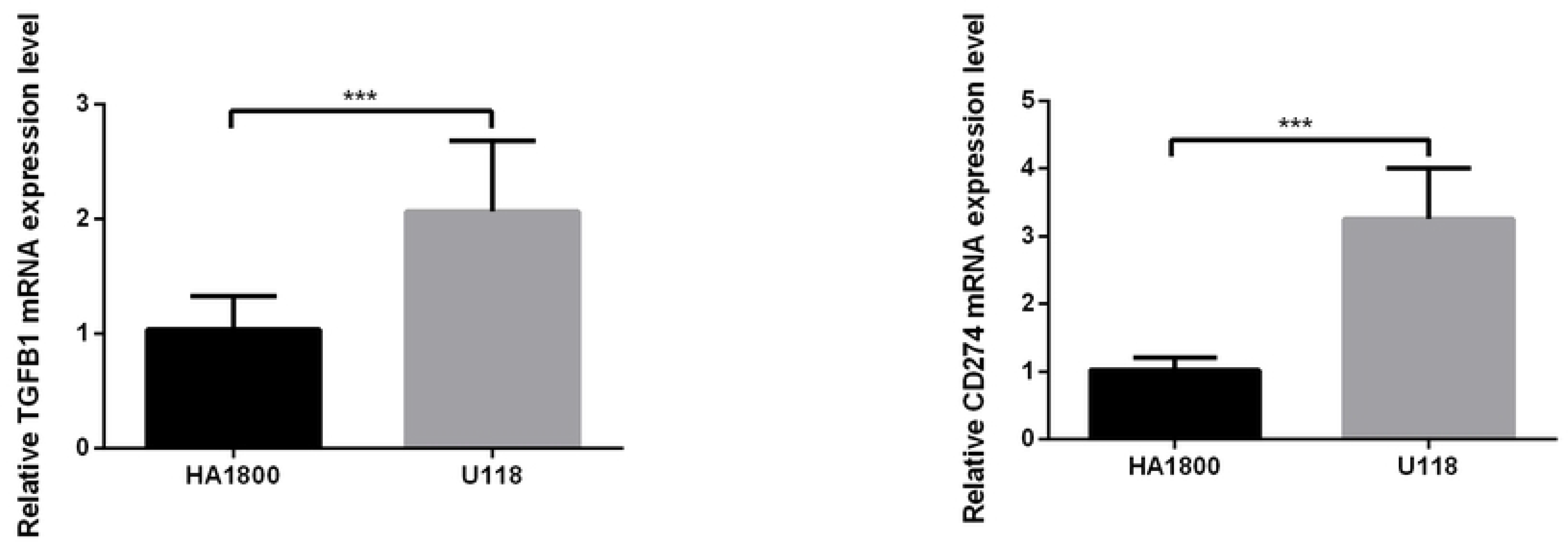
The mRNA expression of TGFB1 and PDL1 in a GBM cell lines and the adjacent cell lines, ***: P < 0.001.

## Discussion

In recent years, the rise of immunotherapy has revolutionized the treatment of GBM, significantly improving the overall survival rate of patients. However, resistance to immune checkpoint inhibitors is manifested in more than half of patients with high PD-L1 expression. ^28^ In addition, immune-related adverse events are common. The current understanding of the mechanisms of resistance to ICB therapy is still limited. An immunosuppressive TME composed of tumor cells, immune cells, and other stromal components may play an important role in ICB resistance.^29^ Therefore, molecular characterization of immunosuppressive TMEs is critical for identifying GBM patients with ICB resistance and thus optimizing immunotherapy strategies.

In our study, the gene expression signals of Tex, immune cells, and stromal components in the TME from GBM were deconvolved using UCC pairs; then, a new class of immunosuppressive GBMs (38.2%) were successfully identified and is defined herein as IDC. Consistent with the depleted immune classes observed in head and neck squamous cell carcinoma and hepatocellular carcinoma,^30–31^ IDCs had higher immune and stromal enrichment scores, indicating the presence of abundant immune cell and stromal components. As expected, IDCs had specific molecular features, including high immune cell infiltration, co-upregulation of multiple inhibitory receptors, enhanced expression of immunosuppressive cytokines, and elevated PD-L1 expression. Among these immune cells, the M2 subtype of tumor-associated macrophages and CD4 Treg cells act as immunosuppressive cells and play an important role in immune evasion and influence ICB therapy.^32–33^ IDCs are widespread across different tumor stages, but have the potential to differ in levels of T-cell exhaustion between early (stage I II) and advanced (stage III) GBM. A previous study showed that severely depleted T cells may undergo apoptosis.^34^ Apoptotic marker gene sets were enriched in late IDC but not early IDC, suggesting that late IDC has higher levels of T cell exhaustion. It also guides clinical medication by predicting the sensitivity of different categories of patients to drugs. We developed a collection of gene-based prediction models by searching for the features of immune depletion classification. These models can accurately forecast the survival prognosis of patients and give patients a solid foundation on which to decide on immunotherapy in a clinic. At the same time, we found that the subtype of immune failure was distributed in the high-risk group, which also verified our previous analysis.

Furthermore, in GBM, the genomic differences (mutation burden and neoantigens) of tumors between IDC and the rest of the classes were also similar to those in other tumors such as head and neck squamous cell carcinoma and hepatocellular carcinoma.38-39 These lines of evidence suggest that tumor-intrinsic mutations may not affect the immunosuppressive microenvironment. The robustness of IDC was successfully validated in three validation datasets. However, this finding requires further validation in GBM patients treated with ICB. Understanding the molecular characteristics of immunosuppressive TMEs is critical for finding successful TEX reversal and immunotherapeutic solutions.

## Conclusion

In conclusion, we identified an immunosuppressive class, accounting for approximately 38.2% of GBM patients, that exhibited potential resistance to ICB therapy and the unique immunosuppressive molecular signature of TME. Our findings provide new insights into understanding the molecular mechanisms of resistance to ICB therapy and tailoring appropriate immunotherapy strategies for patients with different molecular signatures.

## Abbreviations

Immune checkpoint blockade, ICB; Glioblastoma, GBM; Immune depletion class, IDC; central nervous system, CNS; Food and Drug Administration, FDA; programmed cell death protein 1, PD-1; Programmed death-ligand 1, PD-L1; colony-stimulating factor, GM-CSF; Unsupervised Consistent Clustering, UCC; The Cancer Genome Atlas, TCGA; Chinese Glioma Genome Atlas, CGGA; gene set enrichment analysis, ssGSEA; tumor-infiltrating lymphocytes, TILs; Gene ontology, GO; Kyoto Encyclopedia of Genes and Genomes, KEGG; T cell exhaustion, tex; overall survival, OS;

## Declarations

All claims expressed in this article are solely those of the authors and do not necessarily represent those of their affiliated organizations, or those of the publisher, the editors and the reviewers. Any product that may be evaluated in this article, or claim that may be made by its manufacturer, is not guaranteed or endorsed by the publisher.

## Authors’ contributions

Z.T., R.D.designed the study, performed the major data analysis, and drafted the manuscript; Z.T. collected a part of data, performed a part of data analysis, and help to generated fgures. Z.Y. performed a part of data analysis. M.J. helped to generate fgures. W.Z. provided funding source, designed, oversaw, and supervised the project and edited, reviewed, and fnalized the paper. All authors read and approved the fnal manuscript.

## Acknowledgments

The authors thank all the participants for their cooperation.

## Ethics approval

The only data used in this study are the original data of public databases, so no ethical approval is involved

## Data Availability Statement

Datasets analyzed during the current study are available in The Cancer Genome Atlas (TCGA) repository: https://portal.gdc.cancer.gov/, http://www.cgga.org.cn/.

## Consent for publication

Not applicable.

